# Evaluation of sequencing reads at scale using rdeval

**DOI:** 10.1101/2025.02.01.636073

**Authors:** Giulio Formenti, Bonhwang Koo, Marco Sollitto, Jennifer Balacco, Nadolina Brajuka, Richard Burhans, Erick Duarte, Alice M. Giani, Kirsty McCaffrey, Jack A. Medico, Eugene W. Myers, Patrik Smeds, Anton Nekrutenko, Erich D. Jarvis

**Author notes:** Equal contributions (Giulio Formenti, Bonhwang Koo, Marco Sollitto).

## Abstract

**Motivation:** Large sequencing data sets are produced and deposited into public archives at unprecedented rates. The availability of tools that can reliably and efficiently generate and store sequencing read summary statistics has become critical.

**Results:** As part of the effort by the Vertebrate Genomes Project (VGP) to generate high-quality reference genomes at scale, we sought to address the community need for efficient sequencing data evaluation by developing rdeval, a standalone tool to quickly compute and dynamically display sequencing read metrics. Rdeval can either run on the fly or store key sequence data metrics in read ‘sketches’, with dramatic compression gains. Statistics can then be efficiently recalled from sketches for additional processing. Rdeval can convert fa*[.gz] files to and from other popular formats including BAM and CRAM for better compression. Overall, while CRAM achieves the best compression, the gain is marginal, and BAM achieves the best compromise between data compression and accessing speed. Rdeval also generates a detailed visual report with multiple data analytics that can be exported in various formats. We showcase rdeval’s functionalities using human and VGP read data from different sequencing platforms and species. For PacBio long-read sequencing, our analysis shows dramatic improvements both in read length and quality over time, and a benefit of additional coverage for genome assembly.

**Availability and implementation:** Rdeval is implemented in C++ for data processivity and in R for data visualization. Precompiled releases (Linux, MacOS, Windows) and commented source code for rdeval are available under MIT license at https://github.com/vgl-hub/rdeval. Documentation is available using ReadTheDocs (https://rdeval-documentation.readthedocs.io). Rdeval is also available in Bioconda and in Galaxy (https://usegalaxy.org). An automated test workflow ensures the consistency of software updates.

**Supplementary information:** Supplementary data are available at Bioinformatics online.

## Introduction

Over the past twenty-five years, we have witnessed an unprecedented increase in the number of publicly available sequencing data sets (Lewin *et al*., 2022; Larivière *et al*., 2024). Data has been generated using a variety of sequencing platforms, with Illumina short reads being by far the most abundant. Since 2010, Pacific Biosciences (PacBio) and Oxford Nanopore Technologies (ONT) revolutionized sequencing with the introduction of radically new long-read sequencing platforms (Giani *et al*., 2020). Long-read sequencing platforms can routinely generate reads in the order of tens to hundreds of Kbps. While the base calling accuracy of Illumina short reads had been considerably higher than that of long reads for many years, recent improvements in both PacBio and ONT are now rivaling if not exceeding Illumina read quality. In 2022, after 20 years of near-monopoly, the expiration of Illumina short-read patents paved the way to new short-read platforms, including Singular Genomics, Ultima Genomics, MGI, and Element Biosciences’s AVITI (Pollie, 2023).

Sequencing data is made publicly available by large national and international initiatives such as the Sequence Read Archive (SRA) by the US National Center for Biotechnology Information (NCBI, https://www.ncbi.nlm.nih.gov/sra/), the European Nucleotide Archive (ENA, www.ebi.ac.uk/ena/browser/home) by the European Bioinformatics Institute, the DNA Data Bank of Japan (DDBJ, www.ddbj.nig.ac.jp), the China National Genebank (CNGB, https://db.cngb.org), or in project-related repositories such as the GenomeArk (https://www.genomeark.org) by the Vertebrate Genomes Project (VGP) (Rhie *et al*., 2021). Several of these initiatives coordinate through the International Nucleotide Sequence Database Collaboration (INSDC, https://www.insdc.org). Reads are usually stored as compressed archives (e.g. .SRA) or in popular formats, particularly FASTQ. The FASTQ format was developed over two decades ago at the Wellcome Trust Sanger Institute (Cock *et al*., 2010) and later popularized by Illumina to store short-read sequencing data. It reports both sequence and per-base quality information, as well as metadata on the sequencing runs in the headers. As data production globally scales up, and so do storage needs, scientific groups and communities have been testing alternative solutions that can provide better data compression, with or without data loss. Popular options include the SAM/BAM and CRAM formats. The SAM (Sequence Alignment Map) format was originally developed in 2008 as part of the 1,000 Genomes Project to store sequence alignments to a reference (Li *et al*., 2009). SAM files can be losslessly compressed to binary BAM files using BGZF compression, also enabling random lookups through an index file. CRAM was developed in 2011 by the European Bioinformatics Institute (EBI) with the same purpose of compressing SAM records. The CRAM format specification is maintained by the Global Alliance for Genomics and Health (GA4GH) (Rehm *et al*., 2021). CRAM can in principle achieve higher compression by only storing base calls that differ from a reference (Hsi-Yang Fritz *et al*., 2011). Further compression is achieved by storing each SAM column into separate blocks, improving the compression ratio of similar data. CRAM compression can be both lossless and lossy. The latest version, CRAM v3.1 allows additional 7–15% compression via new custom compression codes that rely on bit-packing and other innovations (Bonfield, 2022).

Short-read sequencing generates reads all of the same length, usually within the 100-500 bp range. In short-read sequencing, read length is limited by the sequencing by synthesis (SBS) technology (Giani *et al*., 2020). In long-read sequencing, for both PacBio and ONT, sequencing occurs in real time without DNA amplification, imposing significantly fewer constraints on read length. Initially, PacBio sequencing routinely generated reads above 50 Kbp, with some reads over 100 Kbp. With the advent of the more accurate Circular Consensus Sequencing (CCS) as the standard sequencing approach and the introduction of PacBio High Fidelity (HiFi) reads in 2019 (Wenger *et al*., 2019), read lengths are confined to a narrower 10-20 Kbp range, which provides a good compromise between read length and consensus accuracy. ONT is currently the only technology capable of routinely generating reads over 100 Kbp (ultralong reads; UL) and sometimes above 1 Mbp (whales), but with lower consensus accuracy compared to HiFi.

Read quality is an important factor affecting downstream analyses such as read mapping and genome assembly. Per base quality is self-reported by the sequencing platforms using Phred scores (Q scores, or QV for quality values) (Ewing and Green, 1998). The formula for the Phred score maps the probability of a particular base being incorrect to an integer Q. The most common formula is the Sanger: −10×Log_10_(P). Sanger format can encode a Phred quality score from 0 to 93 using ASCII 33 to 126. In real scenarios, Phred quality score rarely exceeds 60 for reads, but higher scores are possible for individual bases, assemblies or read maps. Phred scores are encoded in FASTQ, SAM/BAM, or CRAM files. Illumina Phred scores rarely exceed Q40, while more accurate short-read platforms such as AVITI can reach Q50. In 2014, to reduce storage footprint, Illumina introduced the concept of binning for Q scores, in which ranges of Q values are assigned the same Q values (Illumina, 2014). By homogenizing the Q values, Q score binning can achieve better data compression as more similar values are easier to compress. Originally, PacBio Continuous Long Reads (CLR) had error rates >10-15% (Q8-10). Estimated Q scores for insertions, deletions, and substitutions were stored in different BAM tags. PacBio read quality improved radically with the adoption of CCS and HiFi (Wenger *et al*., 2019). HiFi reads are CCS filtered for an average quality of at least Q20. HiFi reads usually show average read quality Q27-30. HiFi reads are natively stored in SAM/BAM format and can report significantly higher Q scores (>80) for individual bases in a read. The encoded probability reflects the confidence of a base call against alternatives including substitutions, deletions, and insertions (PacBio BAM format specification — PacBioFileFormats 13.0.0 documentation), and false negative detection of alternative calls can potentially inflate consensus accuracy estimation. In PacBio’s latest sequencing platform, the Revio, Q scores are also binned to increase storage efficiency, and capped at Q40 by default (representing the range [Q40, Q93]). Methods to further improve per-base accuracy are continuously developed, including the popular DeepConsensus approach (Baid *et al*., 2023) that is now integrated in the PacBio sequencing platform, as well as other methods to generate consensus sequences with better calibrated Q scores (Weerakoon *et al*., 2025). ONT reads also had low QV (∼Q10) but have progressively improved over the years. ONT Duplex reads, a special type of library preparation that allows reading both DNA strands that are then merged in a higher-accuracy consensus sequencing, have now quality values capped at Q50 when called using the Dorado base caller, and are also natively stored in SAM/BAM format (Koren *et al*., 2024).

As more sequencing data become available, a single, fast, versatile tool that can compute read summary statistics from a variety of file formats is warranted. As part of gfastar, a tool suite to aid telomere-to-telomere genome assembly, and in the framework of the VGP, which aims to generate high-quality genome assemblies for all vertebrate species, we have developed and present here rdeval (short for “read evaluation tool”). Rdeval can efficiently compute accurate summary statistics, plot read distributions, and convert between different file formats, facilitating the storage and analysis of large data sets.

## Results

Rdeval works by loading sequencing read files, pre-processing them, computing read statistics and optionally saving them in different formats (**Figure 1a**). Rdeval v0.0.5 (the version presented hereinafter) accepts all popular sequence read file formats, including FASTA, FASTQ, SAM/BAM, and CRAM, enabling seamless conversion between these file formats. SAM/BAM and CRAM support is provided by HTSLIB (Bonfield *et al*., 2021). We observed that unaligned BAM and CRAM achieve better compression over plain text and compressed FASTQ files by 6.8-fold and 1.3-fold on average, respectively (**Figure 1b**). CRAM could be considered as the format of choice when high compression efficiency is needed, especially for large repositories, as it achieves the smallest sizes while maintaining essential information, including sequence and base call quality.

**Figure 1.**
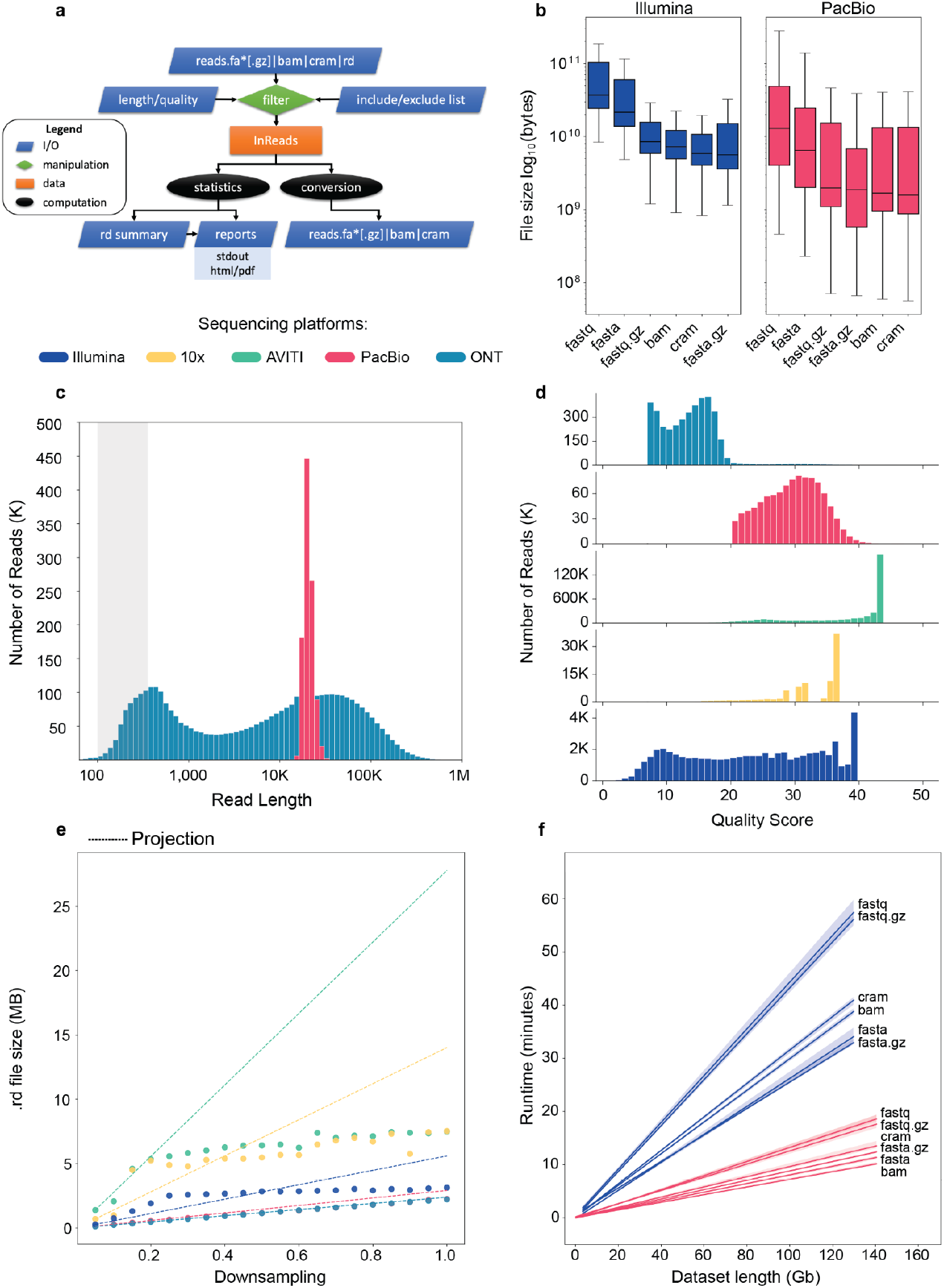
a) Schematic of rdeval workflow. Inputs (blue) include genome assemblies in fasta, fa*[.gz], BAM, CRAM formats and include/exclude lists as bed coordinate files for filtering (green). b) Comparison of compression levels of 84 sequencing data sets included in the VGP project across different file types (FASTA [.GZ], FASTQ [.GZ], BAM, and CRAM) and sequencing platforms (Illumina and PacBio; **Supplementary Table 2**). File sizes are on a logarithmic scale. c) Length distribution of reads from two representative human genomic data sets (CHM13 and HG002). d) Quality distribution for the same data sets. Note that PacBio HiFi reads are at least Q20 and are capped at Q40. e) Relationship between original file size and .rd file size in different data sets: the X-axis represents downsampling, while the Y-axis shows .rd file sizes in MB. The test has been performed on three different sequencing runs from human genomic data sets (**Supplementary Table 3**) generated by different sequencing platforms. The .rd size at the first downsampling level (0.05) has been used to estimate theoretical projections at subsequent steps. Projections are shown in dashed lines. f) Runtime (minutes) versus data set size (Gbp) for 84 sequencing data sets from Illumina (n=37) and PacBio (n=47) platforms across various file formats.

Rdeval can process heterogeneous file formats in the same run, which also allows for generating single sequence archives from multiple files in different formats. Input can optionally be filtered in a pre-processing step to include/exclude sequences based on combinations of criteria. In particular, reads can be filtered by length, e.g. to remove very short reads or to retain only UL reads, and/or they can be filtered by sequence quality to remove low-quality reads or retain only highly accurate reads. Indeed, reads from different state-of-the-art technologies show marked differences in both read length and quality. For instance, in two representative human data sets (**Supplementary Table 3**), short reads are 100-500 bp-long. Illumina data show a wide range of quality scores (Q2-40), with mode at Q40. 10x Genomics linked reads and AVITI reads by Element Biosciences present qualities spanning Q18-43 with high-frequency at Q36 and Q43, respectively. ONT reads show a dispersed bimodal distribution of read lengths, with peaks at 500 bp and 40 Kbp (**Figure 1c**). HiFi reads present an unimodal distribution centered at 20 Kbp with little standard deviation. In terms of quality, ONT reads are mostly below Q20, showing a bimodal distribution centered at Q7 and Q17 (**Figure 1d**). Additionally, a small subset of reads shorter than 1 Kbp close to Q90, possibly due to a base-calling artifact. HiFi reads present quality scores in the range of Q20-40 with the highest frequency at Q30. Rdeval further allows homopolymer compression in linear time without requiring auxiliary space, a feature increasingly useful when dealing with long reads. Interestingly, after homopolymer compression, read data sets are usually on average 29.5% smaller than their uncompressed counterparts (**Supplementary Figure 1**).

Rdeval can compute relevant metrics regarding read size and quality, and these features can be optionally post-processed for downstream analyses including data visualization purposes. Key to this process is the generation of a read ‘sketch’ that stores sorted lengths and Q scores for all reads. The sketch is a binary .rd file, which stores the information needed for downstream analyses. Binary data is further compressed with gzip, such that sketches maintain the integrity of the original data despite achieving remarkable compression. Rdeval also computes md5sum information for the original files using OpenSSL (Ooms) and stores it in the .rd files. Md5sum information can be used to uniquely assign summary statistics to the original files from which they were derived, resolving any ambiguities. Additionally, rdeval can be used to randomly downsample reads through the ‘sample’ option, which allows for the extraction of a specific fraction of reads to create subsampled data sets. This enabled us to investigate the relationship between the original file size and the .rd file size. Compression factors range from 9,000-fold for long-read data sets to 650-fold for short-read data sets on average, essentially removing any overhead when sequence data metrics are needed for data analytics. Our tests show that the .rd file size increases sublinearly with the level of downsampling across all samples, increasing the compression ratio as more data is processed (**Figure 1e**). For instance, in the case of the AVITI data set, we have observed a reduction in file size of approximately 75% when comparing the expected size to the empirical one.

For optimization purposes, rdeval is coded in C/C++, taking full advantage of object-oriented programming. Code is highly commented and a verbose option is available for debugging. Expensive operations are multithreaded to minimize runtime overhead, including file decompression and compression during I/O. Rdeval can compute metrics in O(N) time with runtimes of approximately 5-10 minutes for a typical 30x coverage human data set input using long-read data (PacBio HiFi or ONT), and about 30 minutes for Illumina short reads. Runtime variability depends on both the sequencing platform and the file format. In our experiment with data sets up to 150 Gbp, all runs took less than one hour (**Figure 1f**). Illumina data sets consistently exhibit higher runtimes than PacBio data sets of equivalent sizes, which is due to the need of processing many more reads of shorter lengths. BAM achieves overall runtime best performances for both sequencing platforms while preserving all the information in the data sets. The maximum memory footprint recorded was ∼10 GB, observed in only a few data sets with exceptionally large sizes. On average, however, rdeval has modest RAM requirements (<4 GB). Large outputs are streamed to disk, minimizing memory footprint. Critically, rdeval’s own .rd data format allows extensive parallelization of the process by generating tiny intermediate files that can then be combined into full reports with almost no runtime overhead. This simple procedure is described in rdeval’s ReadTheDocs documentation (https://rdeval-documentation.readthedocs.io). Rdeval sketches format specification is also reported in the documentation.

Compressed data can be used to dynamically display the data sets, generating complex data visualization, including by re-analyzing data without going back to the original data, which is particularly important in the case of large data sets. This is a useful feature, particularly when analyzing several data sets together or in consortia hosting hundreds or thousands of data sets in dedicated repositories. To showcase this functionality, we have computed rdeval sketches for 3,010 individual long-read sequencing runs, i.e. most of VGP’s data with SRA accessions to date. While they represent 63.5 Tb from 4.4 billion sequencing reads (roughly 50 TB if stored as GZIP-compressed FASTQs), the total file size of the sketches is just 9.5 GB. These sketches can be used to analyze the evolution of sequencing technologies over time. For instance, variations in read length can be observed across PacBio VGP data sets generated using different platforms and technologies. The average PacBio read length is 10,803 bp for CLR data generated with a Sequel instrument, 15,599 bp for CLR data produced using the Sequel II platform, 16,116 bp for Sequel II HiFi data, and 12,516 bp for Revio data, respectively (**Figure 2a**). Quality scores have also changed significantly and tend to correlate inversely with read length, though caution should be exercised when interpreting Q scores, distinguishing between actual changes in accuracy and representational changes due to binning, capping, or other factors (**Figure 2b**). We have generated sketch files and sequence reports for all VGP read data in SRA, and the reports are available in GenomeArk (https://genomeark.s3.amazonaws.com/index.html?prefix=downstream_analyses/SRA/). GenomeArk will host rdeval’s sketches and sequence reports for all sequence data made available in the repository moving forward. These sketches are and reports can be useful to investigate the genomic resources. For instance, when we correlated PacBio HiFi coverage to contig N50 in VGP genomes we observed a clear relationship in different clades, suggesting that further increasing coverage will lead to more contiguous assemblies (**Figure 2c**). The correlation is evident in all clades except sharks, which have unusually repetitive genomes for vertebrates, and where clade-specific differences in repeat content may explain the result. Six interesting outliers are noticeable in birds, reptiles (two haplotypes, same species), fish, and mammals (two haplotypes, same species). In birds, the King vulture (*Sarcoramphus papa*) is an outlier with a contig N50 of 43.2 Mbp. The King vulture appears to have particularly large macrochromosomes compared to other birds, which can explain high N50 contig. Similarly, in reptiles the Rock iguana (*Cyclura pinguis*) shows exceptionally high contig N50 values (112.8 Mbp). This is not unexpected owing to the peculiar chromosome structure of the genome, with the three largest chromosomes making up half the genome size and the smallest being ∼300 Mbp. Iguanas have been reported to differ in the accumulation and distribution of interstitial telomeric sequences (ITSs), which suggests chromosomal fusions in some species (Altmanová *et al*., 2016). In fish, the coelacanth (*Latimeria chalumnae*) also shows a high contig N50 (42.0 Mbp), which is likely explained by its unusual genome size (2.9 Gbp) and content compared to non-lobe-finned fish (genome size ∼1 Gbp on average). In mammals, the Botta’s pocket gopher (*Thomomys bottae*) shows very low N50 values (5.9 Mbp for haplotype 1 and 6,2 Mbp for haplotype 2) compared to other mammals despite relatively high sequencing coverage (134,8 Gbp). Genetic studies in Gophers have shown high levels of chromosomal variability across and within populations, with varying chromosome size and numbers, and generally higher than usual repeat content for mammals (Voss *et al*., 2024). This relatively high proportion of repetitive DNA is likely to explain the observed results.

**Figure 2.**
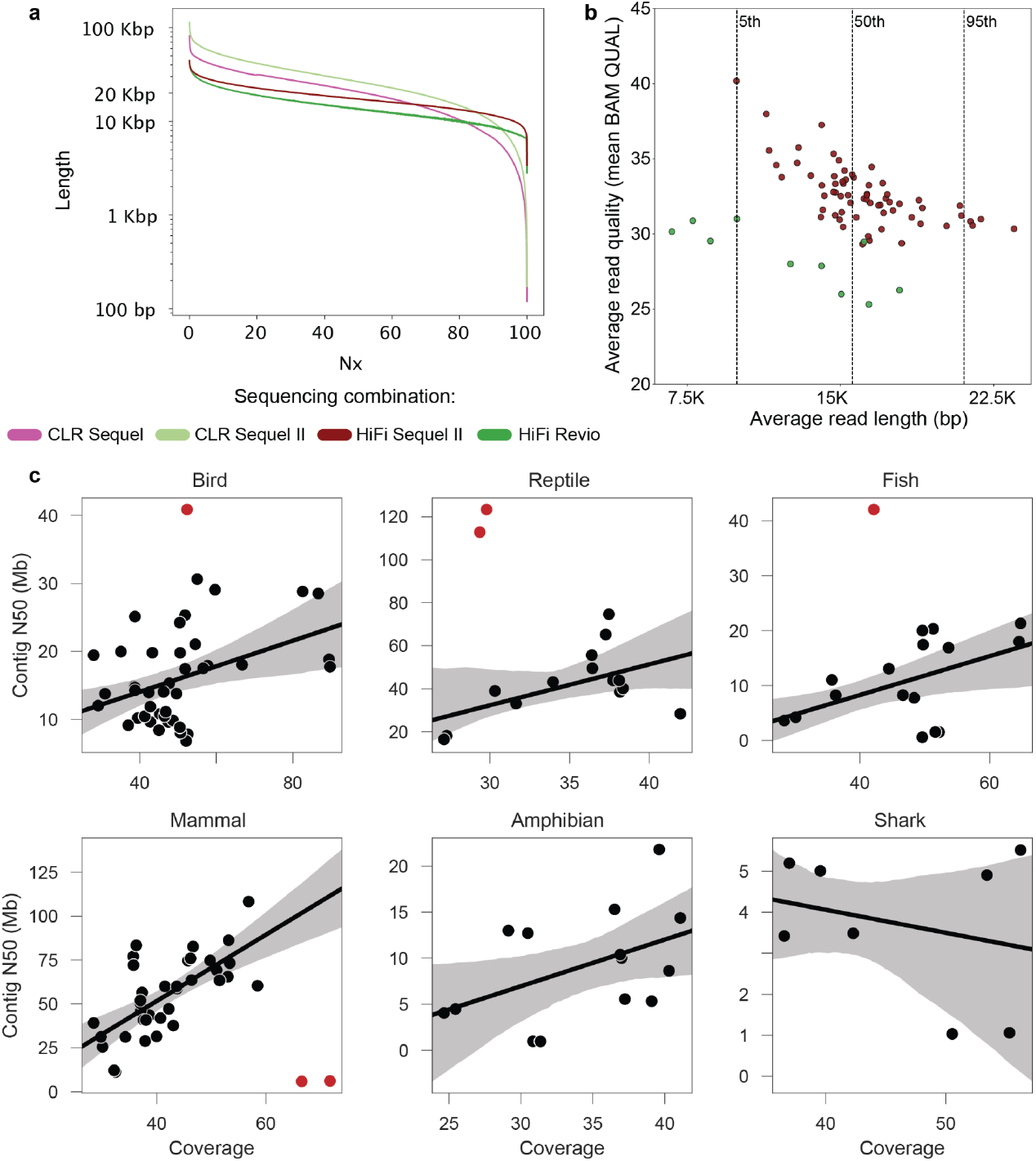
a) Averaged read length Nx plot using all long-read VGP data sets (**Supplementary Table 5**). Most reads were shorter than 100 Kbp on the Sequel Instrument. Read lengths consistently improved with Sequel II. HiFi consensus read lengths were longer on the Sequel II, and shorter with the Revio, likely as a consequence of shorter movie times. b) Average read length vs. average read quality across VGP data sets and instruments. Average read quality is calculated as the mean of the BAM QUAL field. Note that the rq tag in the BAM file represents the estimated “read quality” from Revio and Sequel II, and on Revio, rq may be higher than mean(QUAL) in the final output. Data sets are color-grouped by sequencing platform. Data sets generated by mixed sequencing instruments were excluded from the analysis. Q scores are not available for CLR reads downloaded as FASTQ from SRA. HiFi data shows an inverse relationship between read length and quality, consistent with fewer number of passes available for consensus in longer reads. The difference observed between Sequel II and Revio data is only a representational change, due to capping of Q scores in Revio instruments. c) Correlation between PacBio HiFi sequencing coverage and contig N50 in VGP genomes. Outliers are indicated in red.

Rdeval computes a growing number of general and per-read metrics, including read counts and length, average base quality, base composition, GC content. Many features of rdeval are not available in other tools for sequencing data analysis (**Supplementary Table 1**). Read summaries can be further visualized for one or more files in a HTML report. A representative example of the main graphical outputs is shown in **Figure 3**, with violin plots and histograms illustrating read length, inverse cumulative read length distribution, and the relationship between read length and read quality. See GenomeArk for full report examples.

**Figure 3.**
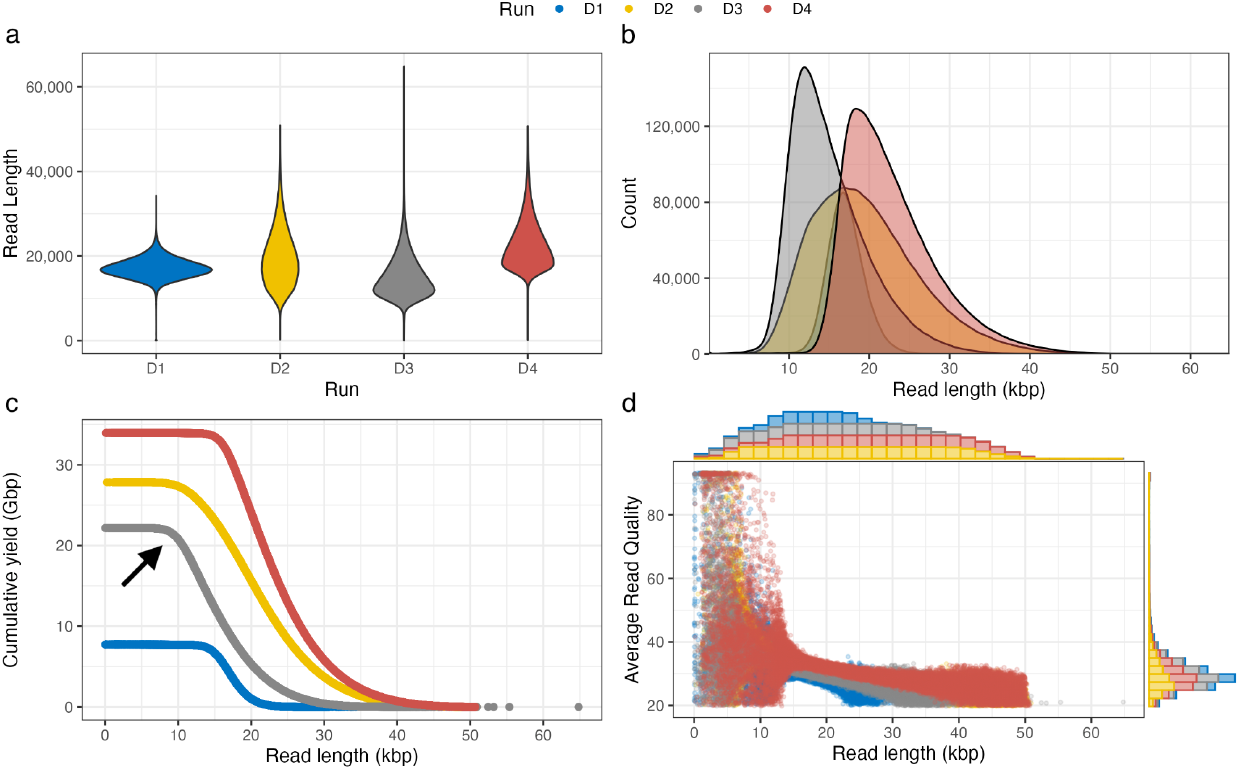
Representative plots from rdeval report using four HiFi sequencing runs from 4 species (D1: *Vipera latastei*, D2: *Amazona ochrocephala*, D3: *Ascaphus truei*, D4: *Gallus gallus*). a) Read length violin plots. D1 shows a tighter read length distribution, and D3 is skewed towards shorter read lengths. b) Read length density plots. Similar to the violin plots, D3 has shorter reads on average and D4 has the longest read lengths. c) Read length inverse cumulative distributions. The distribution can be used to assess read coverage at certain read length cutoffs. For instance, in D3, we can observe that there is about 20 Gbp of coverage for reads 10 Kbp or longer (black arrow). d) Read length vs. Average read quality, plotted with marginal histograms. The plot shows that read length correlates inversely with average read quality at varying magnitudes.

## Discussion and future perspectives

Using rdeval we were able to highlight a number of interesting features in existing read data sets, including the marked differences in read length and quality distribution as a consequence of the rapid evolution of DNA sequencing technologies. For PacBio long-read sequencing, the major transition was from Continuous Long Reads (CLR) generated by the Sequel platform to Sequel II HiFi sequencing. The introduction of the Revio platform coincides with a reduction in average read length. Interestingly, our analysis of homopolymer compression, a widely used approach in genome assembly to overcome the challenges posed by inaccurate base calling of homopolymers in long-read sequencing platforms (Bankevich *et al*., 2022; Rautiainen *et al*., 2023), shows that this strategy results in significant data compression and information loss. Rdeval also showcases how to improve efficiency in large sequencing projects, by choosing more efficient file formats for the storage of raw sequencing reads. BAM and CRAM are valid alternatives to FASTQ, with a slight preference for BAM when a good compromise between compression and data retrieval is needed, particularly when dealing with long reads, whereas CRAM achieves the best compression. Rdeval natively supports these file formats, and is ready to be integrated into production workflows to assess and compare the quality of sequencing runs. It can also become a standard component of large sequencing data projects and repositories to ease access to key read quality metrics. Compared to established tools for sequencing data analysis, rdeval possesses a combination of desirable features not available in similar tools. For instance, FastQC excels at assessing biases within short-read data, such as duplicated sequences and overrepresented k-mers. However, it lacks input manipulation options and generates per-base aggregated metrics incompatible with long-read sequencing. Other popular tools like SeqKit (Shen *et al*., 2016) and seqtk (Li) have additional functionalities (e.g. read length quartiles, CpG counts, etc), but lack support for BAM/SAM available in rdeval and FastQC. Notably, rdeval shows high scalability potential due to multithreaded processing, a feature lacking in FastQC and seqtk. Rdeval offers unique features like CRAM support, coverage calculation, and read sketching in highly compressed binary outputs, which are ideal for analyzing large, mixed-format sequencing data sets. We will continue to add functionalities to rdeval, for example, a ‘quick’ mode that can generate approximate summary statistics by skimming through the data quickly until convergence. We will also evaluate whether to include the functionalities from other tools presented in our comparison table and currently missing from rdeval, to provide all useful sequence manipulation functionalities in a single tool. We will also continue to improve the efficiency of the tool, particularly its runtime and sketch file compression for better performance. Examples of improvements are faster I/O operations and the use of hashing techniques to represent high-frequency reads with identical properties, particularly abundant in short-read data sets, to further increase sketch file compression. Another direction of development will involve testing and integrating more specialized and efficient compression algorithms for genomic sequences such as CoLoRd (Kokot *et al*., 2022) or JARVIS3 (Sousa *et al*., 2024).

## Methods

All runtime tests were run on a server using 4 cores and 30 GB of memory allocated. Plots were generated using 84 different sequencing data sets from the VGP project considering both Illumina (n=37) and PacBio (n=47) sequencing platforms (**Supplementary Table 2**). For Illumina only R1 files were used. The latest CRAM v3.1 was used in the comparisons. To assess quality and read length, downsampled human data sets from CHM13 and HG002 projects were used (**Supplementary Table 3**, AVITI data set can be found at https://s3-us-west-2.amazonaws.com/human-pangenomics/index.html?prefix=T2T/scratch/HG002/sequencing/element/trio/HG002/ins1000/). The same data sets were also used to evaluate the .rd size following downsampling. To allow comparison across technologies, each data set was downsampled to 11 Gbp. The representative report was generated using data from the VGP, in particular PacBio HiFi read files from four species (**Supplementary Table 4**): *Gallus gallus* (SRR30304940), *Vipera latastei* (SRR25383704), *Ascaphus truei* (SRR30223073), *Amazona ochrocephala* (SRR29949495). Files were downloaded from NCBI SRA and converted to FASTQ respectively using the *prefetch* and *fasterq-dump* commands from the SRA toolkit. Binary .rd files were generated for each run to store read length, quality values, and base counts. In order to read the binary .rd files into R, a binary interface was written in R using the *bit64* package. Reports are generated from R Markdown directly from the .rd files using the binary interface. Rdeval sketches for VGP long-read sequencing data were computed by downloading the reads from SRA with *fasterq-dump*. Reads were downloaded and sketches were computed in just two days on a single 32-core machine and only a few hours when using more than 10 nodes in a HPC. Theoretical size of VGP SRA’s as GZIP FASTQs was computed as 2 bytes per base (sequence and quality) * ∼60Tb considering a gzip compression factor of ∼2.5x (headers were ignored). Data visualization is achieved using Rscript. The plots are generated directly from the rdeval .rd files in R using the *ggplot2* and *ggMarginal* packages and the report is rendered using the *render* function from the rmarkdown package. For a thorough analysis of all data sets produced under the VGP umbrella, we examined all SRA sample accessions (ERS, SRS) updated as of January 24, 2025. Some accessions were excluded due to mislabeling or ambiguity regarding whether the data was generated using the Sequel, Sequel II, or Revio platform or cases where sequencing data of a specific biosample were produced using different sequencing instruments (**Supplementary Table 5**). In the coverage vs. contiguity analysis, regression lines were computed excluding outliers. Coverage was computed dividing the total base pair sequences with PacBio HiFi by the assembly size of the primary assembly. All scripts used for figures and analyses in the manuscript are available here: https://github.com/vgl-hub/rdeval-manuscript.

## Author contributions

G.F. implemented rdeval, including the automated test workflows, with contributions from N.B., and A.M.G. and B.K. implemented the data visualization capabilities with contributions from G.F. M.S. compared compression efficiency in different formats. E.D. analyzed the human representative data sets. G.F. and M.S. evaluated rdeval’s performance using the VGP genomes. M.S. B.K. and G.F. analyzed the VGP SRA data sets. J.A.M. and R.B. implemented rdeval in Conda. J.A.M. and P.S. and Galaxy. G.F. and K.M. implemented the documentation on ReadTheDocs. J.B. supported metadata collection and curation. G.F. conceived the study and wrote the manuscript, with contributions from E.D.J. All authors reviewed and approved the manuscript.

## Competing Interests statement

The authors declare no competing interests.

## Acknowledgments

This research is supported by NIH grant number R03HG013362.

---

**Supplementary Table 1:** Comparison of features and operations computed by rdeval against Seqkit (Shen *et al*., 2016), seqtk (Li), and FastQC (Babraham Bioinformatics - FastQC A Quality Control tool for High Throughput Sequence Data). Various metrics, supported formats, and statistics for sequencing data sets are missed in alternative tools.

**Supplementary Table 2:** Sequencing data sets from the VGP project used for testing the runtimes, homopolymer-compression, and level of compression of different formats (FASTA [.GZ], FASTQ [.GZ], BAM, and CRAM).

**Supplementary Table 3:** Representative human sequencing data sets used in the study.

**Supplementary Table 4:** Sequencing data sets from NCBI SRA used to generate representative plots in **Figure 3**.

**Supplementary Table 5:** List of SRA accessions used in **Figure 2a,b**.

**Supplementary Figure 1.**
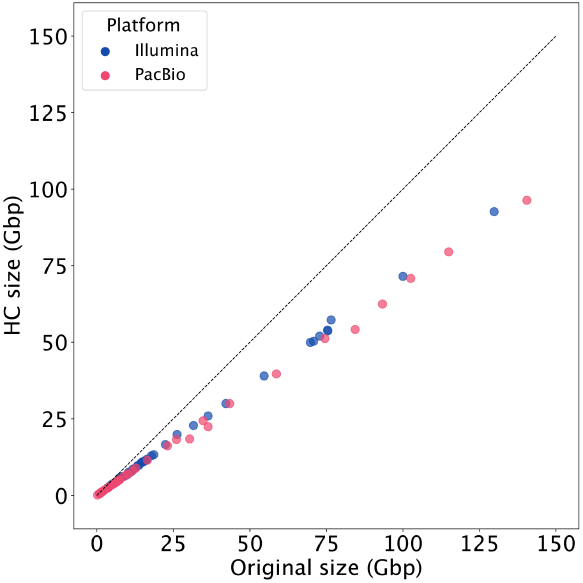
Original file size vs. homopolymer-compressed size for data sets of **Supplementary Table 2**.

## Notes

https://github.com/vgl-hub/rdeval

https://rdeval-documentation.readthedocs.io

### Competing Interest Statement

The authors have declared no competing interest.

### Summary of Updates

- Two figures - Two authors

